# Ultrastructural comparison of dendritic spine morphology preserved with cryo and chemical fixation

**DOI:** 10.1101/2020.03.02.972695

**Authors:** Hiromi Tamada, Jerome Blanc, Natalya Korogod, Carl CH Petersen, Graham W Knott

## Abstract

Previously we showed that cryo fixation of brain tissue gave a truer representation of brain ultrastructure in comparison with a standard chemical fixation method (Korogod et al 2005). Extracellular space matched physiological measurements, there were larger numbers of docked vesicles and less glial coverage of synapses and blood capillaries. Here, using the same preservation approaches we compared the morphology of dendritic spines. We show that the length of the spine and the volume of its head is unchanged, however, the spine neck width is thinner by more than 30 % after cryo fixation. In addition, the weak correlation between spine neck width and head volume seen after chemical fixation was not present in cryo-fixed spines. Our data suggest that spine neck geometry is independent of the spine head volume, with cryo fixation showing enhanced spine head compartmentalization and a higher predicted electrical resistance between spine head and parent dendrite.

## INTRODUCTION

In our previous, parent paper (Korogod et al., 2015), we used a cryo fixation method to study the neuropil of the adult mouse cerebral cortex, comparing it with a standard chemical fixation approach that has been used for many years to preserve the brain for analysis with electron microscopy (eg. Spacek and Harris, 1997; Knott et al., 2002). Korogod et al. (2015) found that cryo fixation appeared to be able to preserve an extracellular space matching various previous *in vivo* measurements, not found in chemically-fixed tissue. Furthermore, synapse density and vesicle docking, as well as astrocytic processes close to synapses and around blood capillaries, appeared to be significantly different between the two fixation methods. This led us in this study to investigate the extent to which dendritic spines might be affected by chemical fixation.

The dendritic spine is a small dendritic protrusion that carries the majority of the excitatory synaptic connections in the adult brain (Colonnier, 1968) with a size and shape that is closely linked with its function (reviewed by: Holtmaat and Svoboda, 2009; Harris and Kater, 1994). The larger the spine head, the larger its synapse (Harris and Stevens, 1988, Arellano et al., 2007; Knott et al., 2006), the number of its receptors (Kharazia et al., 1999; Nogochi et al., 2005; Nusser et al., 1998), and its synaptic strength (Matzusaki et al., 2001). However, the synapse on the spine head is separated from the parent dendrite by the spine neck. This is often thin, compartmentalizing biochemical signals and creating a potentially significant electrical resistance between synapse and dendrite. Changes in the morphology of spine head and neck have been seen after different forms of activity including LTP induction and tetanic stimulation (Lang et al., 2004; Matzusaki et al., 2004, Harvey and Svoboda, 2007; Tønnesen et al., 2014; Fifkova and Anderson, 1981). The understanding of dendritic spine morphology and function has come from many years of analysis using ultrastructural and *in vivo* analysis with electron and light microscopy. However, so far, all the ultrastructural data comes from chemically-fixed samples. These studies show that the brightness of *in vivo* imaged spines closely correlates with their volume measured in serial electron microscopy images (Knott et al., 2006; Cane et al., 2014; Nagerl et al., 2007; El-Boustani et al., 2018). However, the resolution of light microscopy limits any comparison of other features, such as the spine length and neck width. We therefore investigated these parameters in cryo-fixed tissue, using serial section electron microscopy to compare the sizes of spine heads, their neck lengths and neck widths with chemically-fixed tissue. We show that while cryo fixation reveals unchanged spine neck length and head volume, the spine neck is significantly narrower in cryo-fixed tissue compared to chemically-fixed tissue, likely having an important impact upon consideration of the communication between a spine and its parent dendrite.

## MATERIALS AND METHODS

The sample preparation and imaging methods were identical to those used in our previous paper (Korogod et al., 2015) and described briefly below.

### Preparation of cryo-fixed tissue

Adult mice (male, C57BL/6J, 6 - 10 weeks old, N = 5 mice) were deeply anesthetized with isoflurane. After decapitation, the brain was rapidly removed from the skull, and placed on top of a cooled glass Petri dish filled with ice. A small (approximately 2 mm × 2 mm × 0.2 mm piece of cortex was cut from the brain and placed inside the 3 mm diameter and 200 *μ*m cavity of an aluminium sample holder followed by a small drop of 1-hexadecene to remove all the air from around the piece of tissue. This was then frozen using a high-pressure freezer (HPM 100, Leica Microsystems). This entire procedure was completed in less than 90 s from the moment of decapitation.

The frozen samples were transferred to a low temperature embedding device (ASF2; Leica Microsystems) and into 0.1 % tannic acid in acetone for 24 hr at −90° C, followed by 7 h in 2 % osmium tetroxide in acetone at −90°C. The temperature was then raised to −20° C during 14 h, and left at this temperature for 16 hr, and then raised to 4° C during 2.4 hr and left for a further 1 h. Then the liquid was replaced with pure acetone and rinsed at 4°C. Finally, the tissue samples were mixed with increasing concentrations of epoxy resin (Durcupan) and left overnight at room temperature in 90% resin before 100% resin was added the next day for 6 h, and then cured at 60° C for 24 h.

### Preparation of chemically-fixed tissue

Mice (male, C57BL/6J, 6-10 weeks old, N = 3 mice) were anesthetized with an overdose of isoflurane and transcardially perfused with 50 ml of 2.5 % glutaraldehyde and 2.0 % paraformaldehyde in 0.1 M phosphate buffer (pH 7.4). After two hours, the brain was removed and vibratome sliced at a thickness of 80 *μ*m. Slices were then washed thoroughly with cacodylate buffer (0.1 M, pH 7.4), post-fixed for 40 minutes in 1.0 % osmium tetroxide with 1.5 % potassium ferrocyanide, and then 40 minutes in 1.0 % osmium tetroxide alone. They were finally stained for 30 mins in 1 % uranyl acetate in water before being dehydrated through increasing concentrations of alcohol and then embedded in Durcupan ACM (Fluka, Switzerland) resin. The sections were then placed between glass microscope slides coated with mold releasing agent (Glorex, Switzerland) and left to harden for 24 hours in a 65° C oven.

### Electron microscopy imaging

Serial ultrathin sections were cut and collected on copper grids, contrasted with lead citrate and uranyl acetate, and imaged with a transmission electron microscope (TEM) operating at 80 kV (Tecnai Spirit, FEI Company with Eagle CCD camera). Additionally, focused ion beam scanning electron microscopy (FIBSEM) imaging was carried out using the same imaging parameters as described previously (Korogod et al., 2015). This used an imaging voltage of 1.5 kV, and milling voltage of 30 kV and milling current of 80 pA. The pixel size was 5 nm.

### Image processing, analysis, 3D reconstruction, and morphometric measurements

Image series from both the TEM and FIBSEM were registered using the alignment functions in the TrakEM2 plugin of FIJI (Cardona et al., 2012, http://fiji.sc). Spines and parts of the dendrite were segmented with the same software using the *arealist* function. The models were then exported as .obj files into the 3D modelling software (Blender; www.Blender.org). With the *NeuroMorph* tools plugin (Jorstad et al., 2015, 2018; neuromorph.epfl.ch), spine neck cross sectional area, spine neck length, and spine head volume were measured (Figure 1). Spine neck diameter was calculated from the average spine neck cross-sectional area.

**Figure 1.**
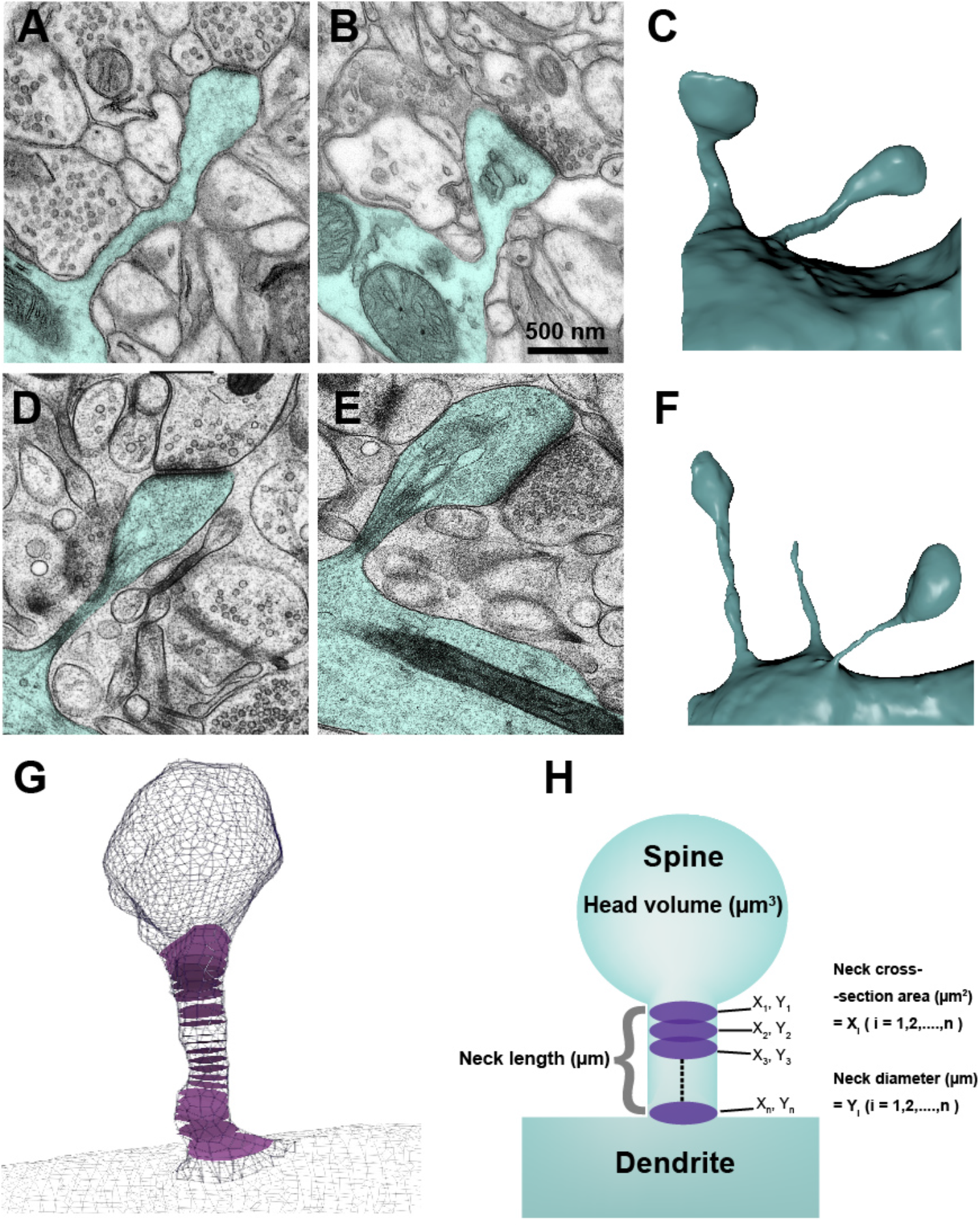
Dendritic spines in chemically fixed (**A**, **B**, **C**) and cryo-fixed (**D**, **E**, **F**) neuropil were reconstructed in 3D from serial electron micrographs. **G**, An example of a 3D model showing cross sections (purple) through the neck region. **H**, From these models the head volume, neck length and cross sectional area were calculated. The neck cross sectional area is an average of several neck cross sectional areas (X_1_, X_2_,…, X_n_) through the spine neck, and the neck diameter (Y_1_, Y_2_,…, Y_n_) was calculated from these values.

### Statistical analysis

All statistical analyses and plots were generated using Graph Pad software Prism 7. The statistical comparisons of chemically-fixed vs cryo-fixed spine neck diameter, cross-section area, length and head volume were performed with unpaired, Kolmogorov-Smirnov tests. Correlation analyses between the quantified parameters were performed with non-parametric Spearman analysis.

## RESULTS

Spines were analysed from 3D models constructed from serial images taken with either TEM (cryo fixation: n = 89 spines, N = 3 mice; chemical fixation: n = 75 spines, N = 1 mouse) or FIBSEM (cryo fixation: n = 27 spines, N = 2 mice; chemical fixation: n = 75 spines, N = 2 mice). No quantitative differences were found between the results obtained by TEM and FIBSEM, and the results were therefore pooled. The tissue was prepared with identical protocols to those used in the parent paper (Korogod et al., 2015). As before, these revealed very little extracellular space with chemical fixation (Figure 1), while cryo fixation showed elements more dispersed with significant spaces separating them. The cryo-fixed membranes also had a distinct appearance from those exposed to chemical fixative. Cryo-fixed membranes were smoothly curving and lacked wrinkles typically seen after chemical fixation. Furthermore, the cryo-fixed tissue showed many thin processes with diameters as narrow as 100 nm (Figure 1).

Quantifying spine geometry, we found that the spine neck cross sectional area was significantly narrower in the cryo-fixed samples (cryo fixation, 0.0159 ± 0.0185 *μ*m^2^; chemical fixation, 0.0284 ± 0.0197 *μ*m^2^; unpaired Kolmogorov-Smirnov test, p < 0.0001; Figure 2A). The mean spine neck diameter was calculated from the measurements of spine neck cross sectional area and found to have a mean diameter of 128 nm in cryo-fixed tissue compared to 182 nm in chemically-fixed tissue (cryo fixation, 128.0 ± 62.8 nm; chemical fixation, 182.4 ± 54.5 nm; unpaired Kolmogorov-Smirnov test, p < 0.0001; Figure 2B). Spine length measurements showed a large diversity in each group, but no statistical difference between the two fixation conditions (cryo fixation, 0.770 ± 0.550 *μ*m; chemical fixation 0.836 ± 0.569 *μ*m; unpaired Kolmogorov-Smirnov test, p = 0.53) (Figure 3C). Measurements of the spine head volume also showed no difference between chemically-fixed and cryo-fixed tissue (cryo fixation, 0.069 ± 0.062 *μ*m^3^; chemical fixation, 0.081 ± 0.087 *μ*m^3^; unpaired Kolmogorov-Smirnov test, p = 0.65) (Figure 3D).

**Figure 2.**
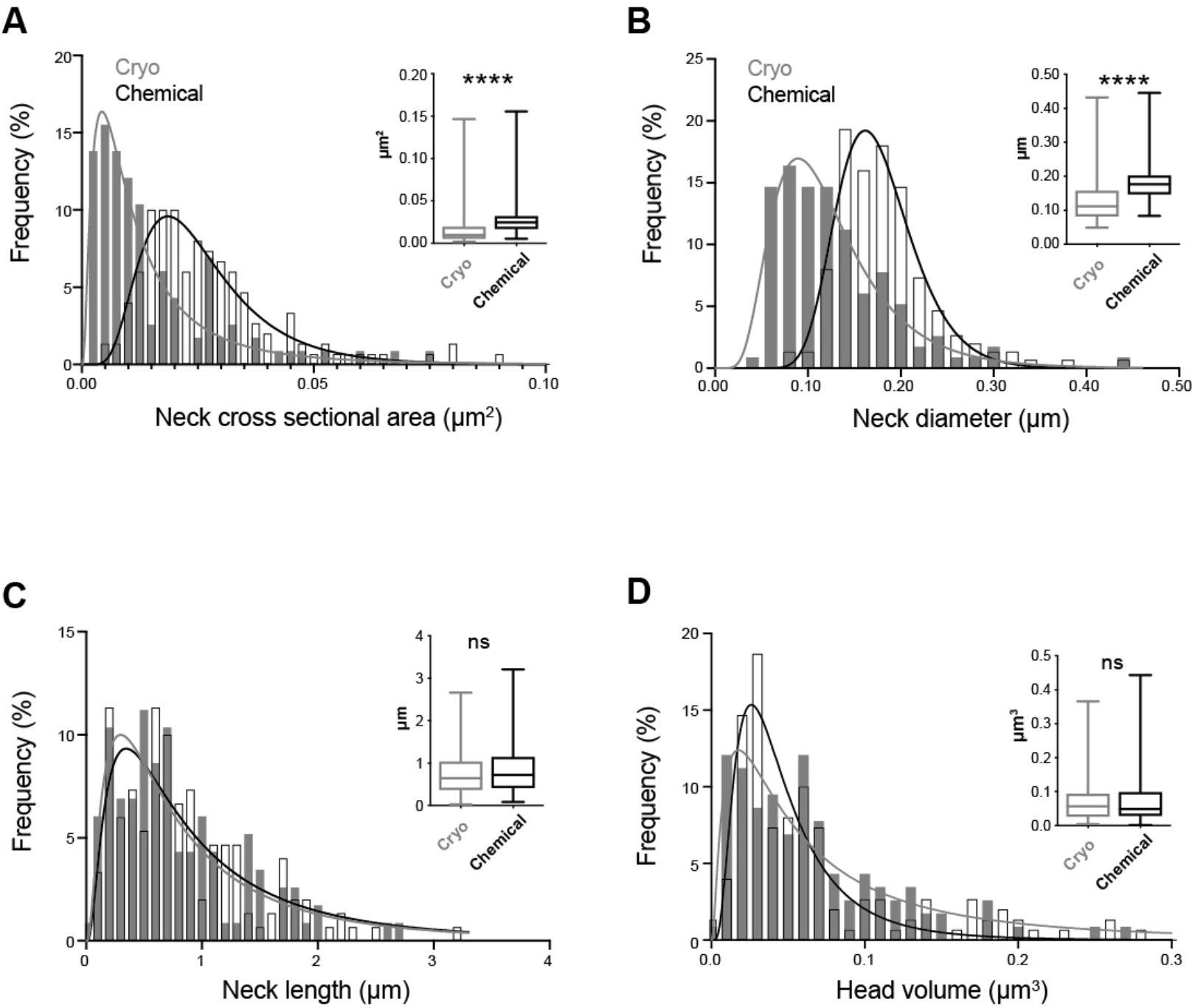
Cryo-fixed dendritic spines show reduced neck widths. **A**, Distribution of neck cross sectional area measurement in cryo-fixed (n = 116 spines, N = 5) and chemically-fixed (n = 150, N = 3) cortex (p < 0.0001, Kolmogorov-Smirnov). Curves show log-normal fits (R^2^ values of 0.93 for cryo-fixed spines; 0.94 for chemically fixed). Inset shows a box plot indicating median, interquartile range and minimum and maximum values. **B,** The distribution of neck diameter calculated from the cross-sectional area measurements in cryo-fixed and chemically fixed cortex (p < 0.0001, Kolmogorov-Smirnov). Log-normal curves have R^2^ values of 0.95 for cryo-fixed spines and 0.97 for chemically fixed. **c**, The distribution of neck length measurements in cryo-fixed and chemically-fixed cortex (p = 0.53, Kolmogorov-Smirnov). Log-normal curves have R^2^ values of 0.80 for cryo-fixed spines and 0.76 for chemically fixed. **C**, Distribution of head volume measurement in cryo-fixed and chemically-fixed cortex (p = 0.56, Kolmogorov-Smirnov). Log-normal curves have R^2^ values of 0.89 for cryo-fixed spines and 0.86 for chemically-fixed.

**Figure 3.**
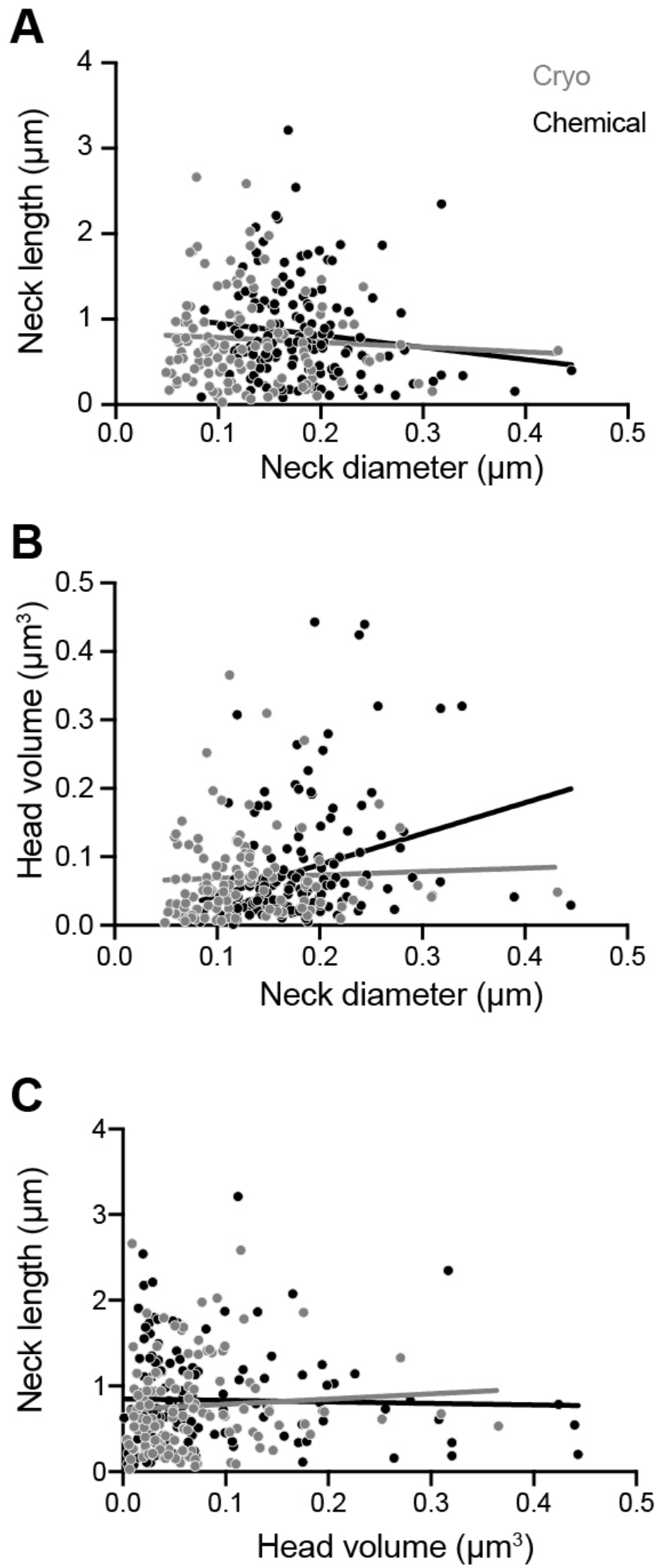
Chemically-fixed, but not cryo-fixed, spines show a correlation between neck width and head volume. **A**, Correlation between neck diameter and neck length for cryo-fixed spines (Spearman r= −0.020, p = 0.83, n = 116 spines, N = 5 mice) and chemically-fixed spines (Spearman r = −0.156, p = 0.054, n = 150, N = 3 mice). **B**, Correlation between neck diameter and head volume; cryo fixed; r= 0.145, p = 0.121; chemically fixed; r = 0.358, p < 0.0001, Spearman). **C**, Correlation between head volume and neck length; cryo fixed; r = 0.124, p = 0.184, chemically fixed; r = 0.0441, p = 0.592.

Previously, we found that the size of synapses was not different comparing the two fixation conditions (Korogod et al., 2015). Here, we further report that synapse area correlates with the volume of the spine head in cryo-fixed tissue (R^2^ = 0.73, n = 33 spines) (Supplementary Figure 1), similar to previous reports studying chemically-fixed tissue (reviewed by Bourne and Harris, 2008).

**Supplementary Figure 1.**
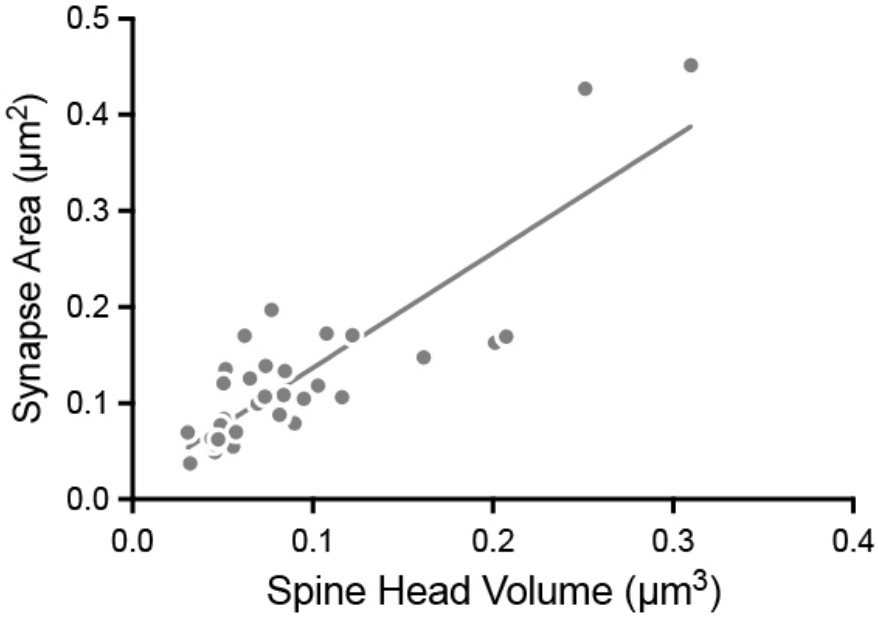
Dendritic spine head volume correlates with the size of the synapse in cryo-fixed neuropil. Plot of synapse area against spine head volume for 33 spines (R^2^ = 0.73).

We next examined correlations between spine head volume, spine neck diameter and spine neck length (Figure 3). Similar to a previous study (Arellano et al., 2007), we found a weak correlation in the chemically fixed spines between head volume and neck diameter (r = 0.36; p < 0.0001; non-parametric Spearman test). This was not observed, however, in the cryo-fixed spines (r = 0.14; p = 0.12; non-parametric Spearman test). We also found no correlation between the head volume and neck length in either cryo-fixed or chemically fixed spines (cryo fixation, r = 0.124, p = 0.18; chemical fixation, r = 0.044, p = 0.59, non-parametric Spearman test) and also no correlation between the neck length and the neck diameter (cryo fixation, r = −0.020, p = 0.83; chemical fixation, r = −0.156, p = 0.054, non-parametric Spearman test).

The correlation of spine head volume and spine neck diameter in chemically-fixed tissue could be due to significant structural differences in the spine neck comparing between larger and smaller spines. Previous EM analyses have shown that spines with larger heads tend to be those where membranous structures, such as endoplasmic reticulum (ER), are present (Spacek and Harris, 1997). If, by their presence, these structures enlarge the neck then this may have an influence on the correlation between head volume and neck thickness. We therefore investigated this question by grouping dendritic spines (chemical fixation, n = 75; cryo fix, n = 98) according to whether or not membranes were present in the neck (Supplementary Figure 2). Though we scored this as ER, it is possible that they could be cisterns or vesicles. We could not classify these unequivocally in the cryo-fixed material because of the lower contrast staining of the cryo-fixed membranes, and in many cases the very thin spine necks would have required a higher resolution method, such as electron tomography to visualize the neck contents more clearly (Supplementary Figure 2A). This analysis showed that spine neck length and the spine neck diameter were not different if membranes were present or absent, neither in cryo-fixed tissue nor in chemically-fixed tissue (neck length, chemical fixation p = 0.95, cryo fixation p = 0.99; spine neck diameter, chemical fixation p = 0.07, cryo fixation p = 0.81; Supplementary Figure 2B, C). Therefore, the presence of intracellular compartments within the neck do not appear to determine its thickness. However, spines in which ER is present in the spine neck do have larger heads after both cryo and chemical fixation (chemical fixation p < 0.0001, cryo fixation p = 0.0074; Tukey’s multiple comparison test; Supplementary Figure 2D), in agreement with a previous study of chemically-fixed spines (Spacek and Harris, 1997).

**Supplementary Figure 2.**
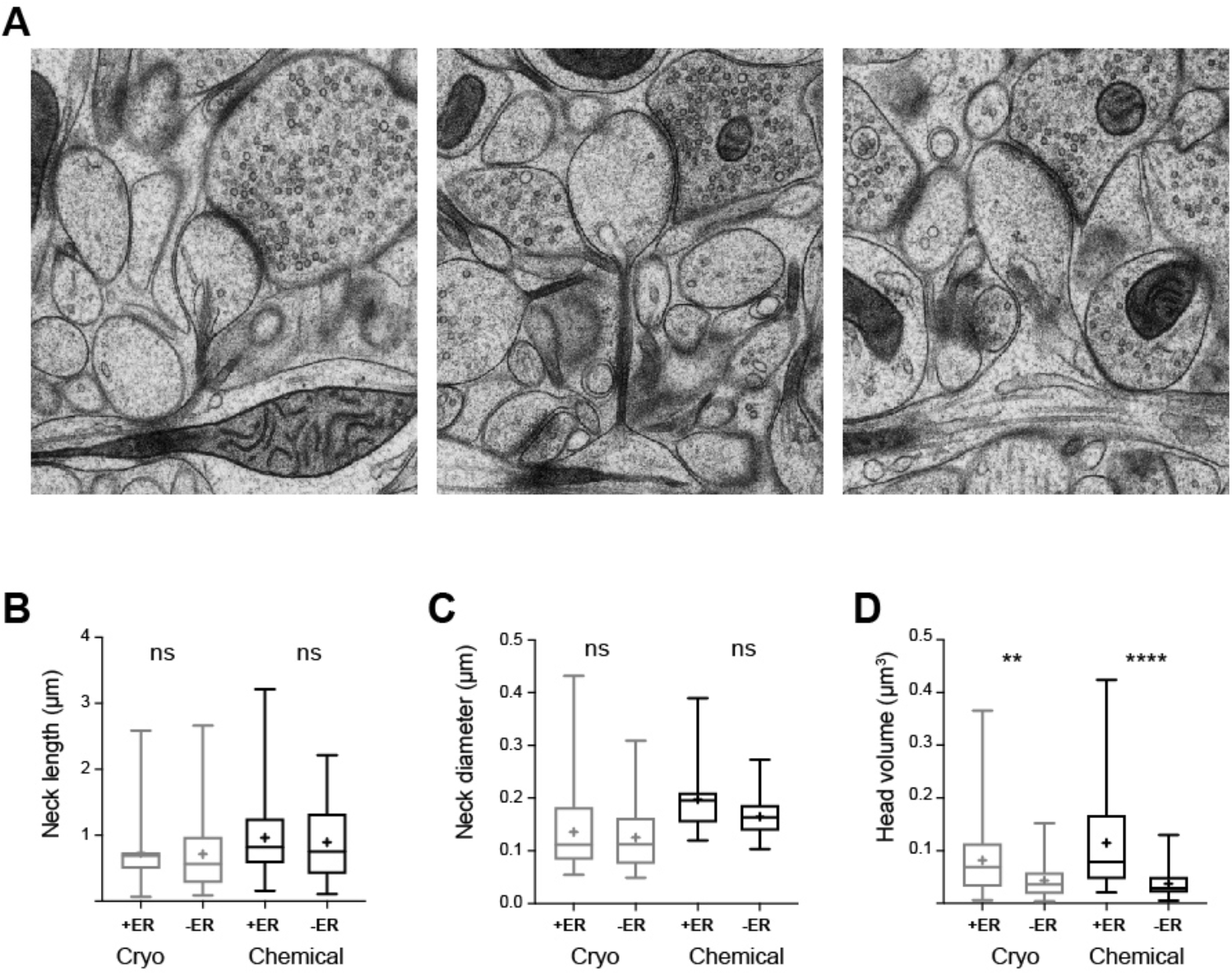
Dendritic spine geometry and the presence of membranes in the spine neck. **A**, Spine neck length and, **B**, spine neck diameter were not significantly different between spines with (+ER) and those without (−ER) membranes in the neck (spine neck length: cryo-fixed p = 0.99, chemical fixation p = 0.95; spine diameter: cryo fixation p = 0.81, chemical fixation, p = 0.08; Tukey’s multiple comparison n = 98 cryo-fixed spines, n = 75 chemically-fixed spines). **C**, Spine head volume was larger in spines with membranes in the neck (cryo fixation p = 0.007; chemical fixation p < 0.0001; Tukey’s multiple comparison). Box plots showing median, interquartile range, minimum and maximum values, and mean (cross). Each group is divided according to whether or not membranes were present in the spine neck (+ER, or −ER).

Finally, we considered how the electrical resistance of the spines would change on the basis of the different morphological parameters measured (Figure 4). We used Ohm’s law and the formula R = ρ × spine neck length / spine neck cross sectional area, with a resistivity constant ρ = 109 Ω cm (Cartailler et al., 2018). The mean value of the resistance of spines in cryo-fixed cortex was 109.7 ± 113.4 MΩ (spine number n = 116, N = 5), and in chemically-fixed cortex was 44.1 ± 37.8 MΩ (spine number n = 150, N = 3) showing a significant difference in their distributions (p < 0.0001, unpaired, Kolmogorov-Smirnov test; Figure 4A). We found obvious correlations between resistance and spine neck cross sectional area (cryo, r = −0.74, p < 0.0001, n = 116 spines, N = 5 mice; chemical, r = −0.64, p <0.0001, n = 150 spines, N = 3 mice, correlation, non-parametric Spearman; Figure 4B). Spine resistance and spine neck length were also clearly correlated (cryo, r = 0.65, p < 0.0001; chemical, r = 0.84, p < 0.0001; Figure 4C). However, we did not find any strong relationship between the resistance of the spine neck and its head volume in either the cryo-fixed or chemical-fixed cortex (cryo, r = −0.014, p = 0.88; chemical, r = −0.16, p = 0.049; Figure 4D).

**Figure 4.**
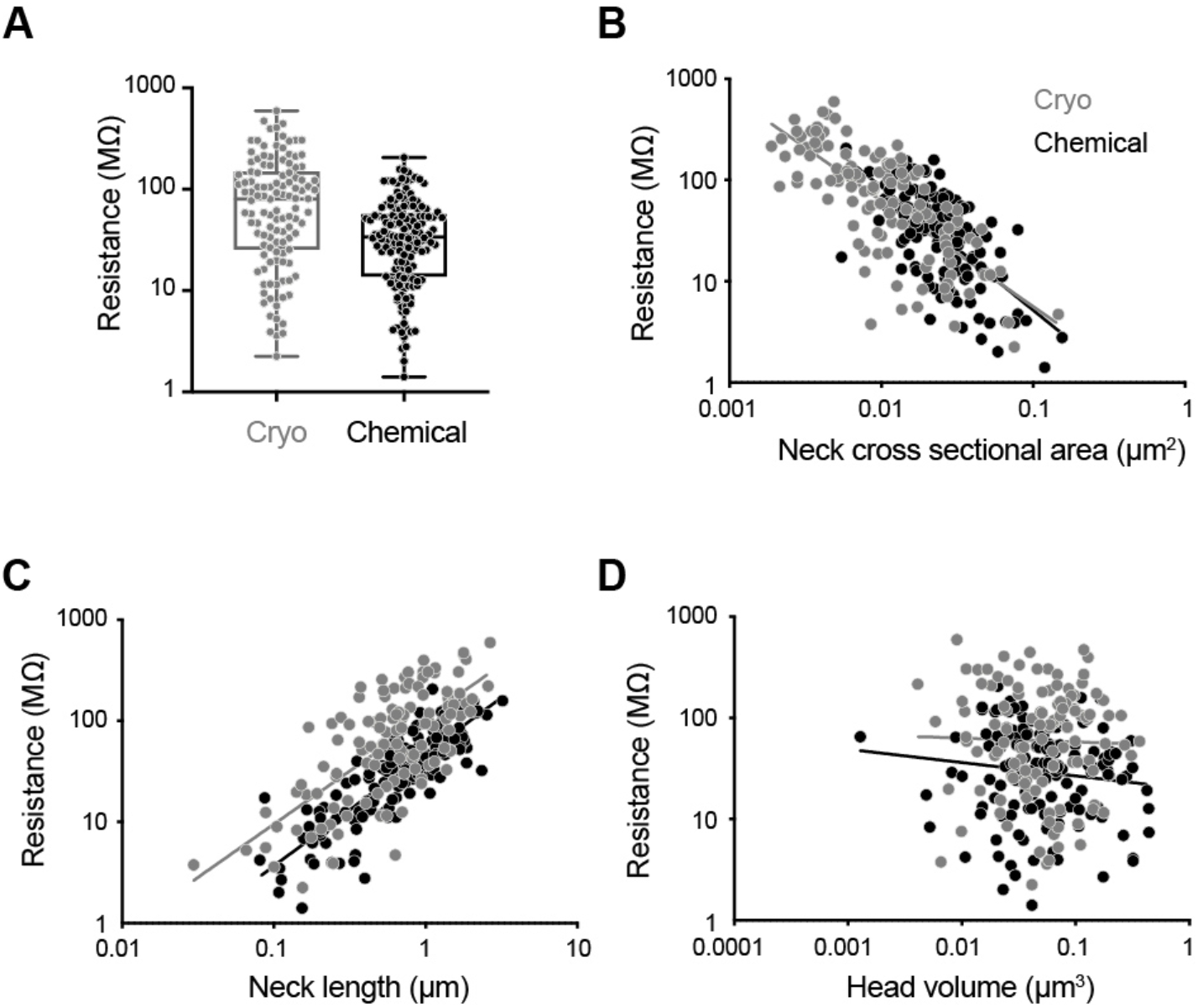
Computed spine neck resistances are higher in cryo-fixed cortex compared to chemically-fixed cortex. **A**, Dendritic spine neck resistances were calculated with resistivity constant of 109 (Ω cm) for cryo-fixed and chemically fixed cortex (cryo-fixed spines n = 116 spines, N = 5 mice; chemically fixed spines n = 150 spines, N = 3 mice). Cryo-fixed spines have higher resistance, graph showing median, range and distribution of resistances (p < 0.0001, unpaired, Kolmogorov-Smirnov test). **B**, Correlation between neck cross sectional area and resistance (cryo fixed, r = −0.79, p < 0.0001; chemical fixation, r = −0.64, p < 0.0001). **C**, Correlation between neck length and resistance (cryo fixed, r = 0.66, p < 0.0001; chemical fixation, r = −0.83, p < 0.0001). **D**, Correlation between head volume and resistance (cryo; r= −0.073, p = 0.43, chemically fixed; r = −0.16, p = 0.049).

## DISCUSSION

The aim of this study was to gain a better understanding of the native ultrastructure of dendritic spines, and to investigate if chemical fixation alters the morphology of dendritic spines, a neuronal structure that for many years has been the focus of considerable research due to its significant role in neuronal connectivity and plasticity (Harris and Kater, 1994; Holtmaat and Svoboda, 2009). We found that the volume of spine heads and the length of spine necks are unaltered by chemical fixation, and only the size of the spine neck is changed, being 42 % thicker in diameter in chemically-fixed tissue compared to cryo-fixed tissue. The thinner spine necks observed in cryo-fixed tissue suggests a larger electrical resistance coupling spines to parent dendrites than previously indicated from analysis of chemically-fixed tissue.

Many quantitative aspects of dendritic spine structure appear to remain unchanged comparing chemically-fixed and cryo-fixed tissue, and are in agreement with previously published work. Spine head volume in cryo-fixed tissue was 0.069 *μ*m^3^, which was not different to the spine volume in chemically-fixed tissue of 0.081 *μ*m^3^ (Figure 2). Our spine volume measurements are similar to other studies in the same region of the mouse brain (0.06 ± 0.04 *μ*m^3^, layer 4 (Rodriguez-Moreno et al., 2017); 0.075 ± 0.005 *μ*m^3^ - 0.079 ± 0.006 *μ*m^3^, layer 1, adult and aged mice, layer 1(Calì et al., 2018). Similarly, our measurement of mean spine length after cryo-fixation of 0.77 ± 0.55 *μ*m was not different from the length of 0.84 ± 0.57 *μ*m in chemically-fixed tissue (Figure 2), and is similar to other studies, for example rat CA1 hippocampus, 0.45 ± 0.29 *μ*m (Harris and Stevens, 1989); rat cerebellum, 0.66 ± 0.32 *μ*m (Harris and Stevens, 1988); and mouse layer 2/3 pyramidal cells, 0.66 ± 0.37 *μ*m (Arellano et al., 2007). Despite the fact that chemical fixation causes important changes to the extracellular space and astrocytic arrangement (Korogod et al., 2015), the overall structure of dendritic spines appears to be resilient to the alterations that the aldehydes are causing.

On the other hand, our results reveal that spine neck thickness is strongly affected by chemical fixation, with a mean diameter of 128 nm in cryo-fixed tissue compared to 182 nm in chemically-fixed tissue (Figure 2). The finding that spines are narrower in the cryo-fixed tissue raises the question as to what could be happening during the conventional aldehyde preservation method that would explain their widening. Spines are highly dynamic structures, with their shape changing during development and also adulthood (Nimchinsky, 2002), and in concert with changes in synapse size (Sala and Segal, 2014; Knott et al., 2006). To explain dynamic spine morphology, we need to consider the structural components. Actin is present throughout spines, with branched and linear actin being present in different proportions in the neck and head (Matus, 2000, Korobova and Svitkina, 2010). This cytoskeleton determines the size and shape of the spine, with synaptic activation capable of altering the quantities of filamentous actin, and changing spine shape (Honkura et al., 2008). Modelling of dendritic spines using the physical properties of its different structural components suggest the presence and arrangement of different proteins, such as actin in the neck, maintain its diameter (Miermans et al., 2017). This is supported by evidence of a clear periodic arrangement of actin rings in the spine neck (Bär et al., 2016) as well as various experiments showing that disruption of the actin regulatory pathways lead to changes in the spine shape (Basu and Lamprecht, 2018). However, preserving the integrity of actin is difficult, due to its sensitivity to aldehyde fixatives such as the ones used here (Boyles et al., 1985). Therefore, if the shape of spines is critically dependent on the arrangement of actin, which is dismantled on exposure to aldehydes, it is perhaps not surprising that spine necks are not the same in cryo-fixed and chemically-fixed tissue.

Some spine necks have also been shown to be rich in other proteins, such as the GTPases from the septin family (Tada et al., 2007), thought to be involved in the restriction of movement of certain membrane proteins out of the spine head (Ewers et al., 2014). A differential protein concentration within this narrowed part of the spine could exert on oncotic pressure pulling water into this region during chemical fixation.

Our data furthermore reinforce the results from light microscopy showing that the morphology of the spine neck is independent of the spine head volume (Araya et al., 2014; Tønnesen et al., 2014). Previous electron microscopy analysis had suggested a correlation, though only a weak one, between these parameters (Arellano et al., 2007; Bartol et al., 2015), which we also show here with chemical fixation. However, cryo-fixed spines, presumed to be in their native state, show no correlation. Spine neck thickness, therefore, appears to be independent of the spine head volume, and therefore synapse size.

Functionally, the smaller spine widths observed in cryo-fixed tissue without change in spine neck length suggest that spine neck resistances might be higher than that implied by previous studies of chemically-fixed tissue. Assuming cytosolic resistivity of 109 Ω cm (Cartailler et al., 2018), we estimate a median spine neck resistance of 80 MΩ ranging from 2 MΩ to 595 MΩ in cryo-fixed tissue compared to 34 MΩ ranging from 1 MΩ to 206 MΩ in chemically fixed tissue. It is important to note that these are likely lower-bound estimates, since the presence of intracellular organelles in the spine neck will further increase spine neck resistance. Our results are consistent with estimates from physiological analyses (eg. Bloodgood & Sabatini, 2005; Grunditz et al., 2008). At the upper end of the range of spine neck resistance, there could be a considerable impact upon spine voltage dynamics during synaptic signaling, with a 1 GΩ spine neck resistance causing a 1 mV drop in potential from spine to parent dendrite for each 1 pA of synaptic current. Given that synaptic currents are typically considered to be in the range of tens of pA, our data are consistent with tens of mV difference between spine head and dendrite for the smallest spine neck diameters. For these spines, regulation of spine neck diameter could therefore contribute to functional synaptic plasticity as previously proposed (Bloodgood and Sabatini, 2005; Grunditz et al., 2008). Our data showing smaller spine neck width in cryo-fixed tissue therefore suggest that spine neck resistance may play a more important role in synaptic signaling compared to estimates based on chemically-fixed tissue.

## Acknowledgements

We thank Lucie Navratilova, at the Center of Electron Microscopy EPFL, for help with the FIBSEM imaging; Catharine Maclachlan, Stéphanie Clerc, and Marie Crosier at BioEM for help with sample preparation.

